# Inferring the demographic history of tetraploid species from genomic data

**DOI:** 10.1101/2021.07.10.451876

**Authors:** Camille Roux, Xavier Vekemans, John Pannell

## Abstract

Genomic patterns of diversity and divergence are impacted by certain life history traits, reproductive systems and demographic history. The latter is characterised by fluctuations in population sizes over time, as well as by temporal patterns of introgression. For a given organism, identifying a demographic history that deviates from the standard neutral model allows a better understanding of its evolution, but also helps to reduce the risk of false positives when screening for molecular targets of natural selection. Tetraploid organisms and beyond have demographic histories that are complicated by the mode of polyploidisation, the mode of inheritance and different scenarios of gene flow between subgenomes and diploid parental species. Here we provide guidelines for experimenters wishing to address these issues through a flexible statistical framework: approximate Bayesian computation (ABC). The emphasis is on the general philosophy of the approach to encourage future users to exploit the enormous flexibility of ABC beyond the limitations imposed by generalist data analysis pipelines.

## 1 Introduction

The living world is characterised by an organisation of individuals into different discrete groups, partially or totally reproductively isolated from each other: species. Within a given species, genetic information at a given locus may be either invariant, i.e. shared by all individuals of that species, or polymorphic, i.e. with different segregating alleles. The relative importance of such monomorphic and polymorphic loci along genomes differs between species and can be empirically quantified by analyzing whole-genome sequences of individual samples of a given species. The theoretical framework offered by population genetics seeks to explain the origins of molecular diversity segregating within species, but also to understand how evolutionary forces act over time on this genetic variation: either to maintain it or to decrease it. Thus, with varying degrees of consequences, mutation, migration, genetic drift and different forms of natural selection will both impact the proportion of polymorphic positions in genomes and allele frequencies within populations. The recent acquisition of large-scale population genomic data for non-model species has facilitated investigations into the relative roles of these different forces in genome evolution [1, 2]. In addition to the perspectives of bringing answers to fundamental evolutionary questions offered by the recent sequencing technologies, they also pave the way for numerous applications in conservation biology. Indeed, assuming that the species is a fundamental unit in conservation biology programs, high-throughput sequencing methods [3, 4] are now greatly facilitating the task of attributing individuals to different biological groups. In a complementary manner, the identification of genomic regions involved in local adaptation is necessary to define relevant ecotypes for reintroduction management project of endangered populations [5]. A central point of this chapter is to highlight the general importance of characterising the demographic context in which species have evolved: how have effective population sizes fluctuated over time? How important is migration in the genetic composition of the population? Importantly, for methodological reasons since many of the inferential biases associated with downstream sequencing analyses are likely to arise when demography is neglected. This can be illustrated by current practices in population genomics. A popular way to identify genomic regions that have recently experienced spatially-divergent selection is based on the “genomic scan” of the *F*_ST_ statistic, i.e. a survey of variation of this statistic across genomic regions aiming to identify regions with higher or lower values than the genomic background. *F*_ST_ quantifies the variance of allele frequencies among populations [6]. The aim of these genomic scans for *F*_ST_ outliers is then to identify *F*_ST_ values that cannot be explained by a drift-mutation-migration equilibrium alone. The usual interpretation of an apparent increased variance is the action of strong and local adaptive processes leading to contrasted directional selection effects in different populations. And on the other tail of the *F*_ST_ genomic distribution, an apparently reduced variance in allele frequencies is interpreted as the action of balancing selection or a local increase in introgression rate. Such *F*_ST_ outliers methods have emerged early in molecular population genetics, before the discovery of low or high-throughput nucleotide sequencing, in order to infer the expected *F*_ST_ distribution under the assumption of selective neutrality, and then, to test for loci that are outliers of such a null distribution [7]. However, numerous problems associated with this approach were raised very shortly after its publication, notably pointing to the large influence of migration patterns and population size on the theoretical variance of *F*_ST_ that cannot be so easily neglected, regardless of the ploidy level [8, 9]. While Lewontin and Krakauer’s seminal paper has gained impact in the modern literature with the advance of high-throughput sequencing methods [10], the contemporary caveats associated with it have not received the same renewed attention [11], indicating that the importance of demographic inferences beyond their descriptive aspects should be recalled. As illustrated by Lotterhos and Whitlock by simulating the neutral evolution of metapopulations placed in different landscapes, alternative demographic histories can lead to different null distributions of *F*_ST_ sharing the same genomic average (Fig. 1; [12]).

**Fig. 1.**
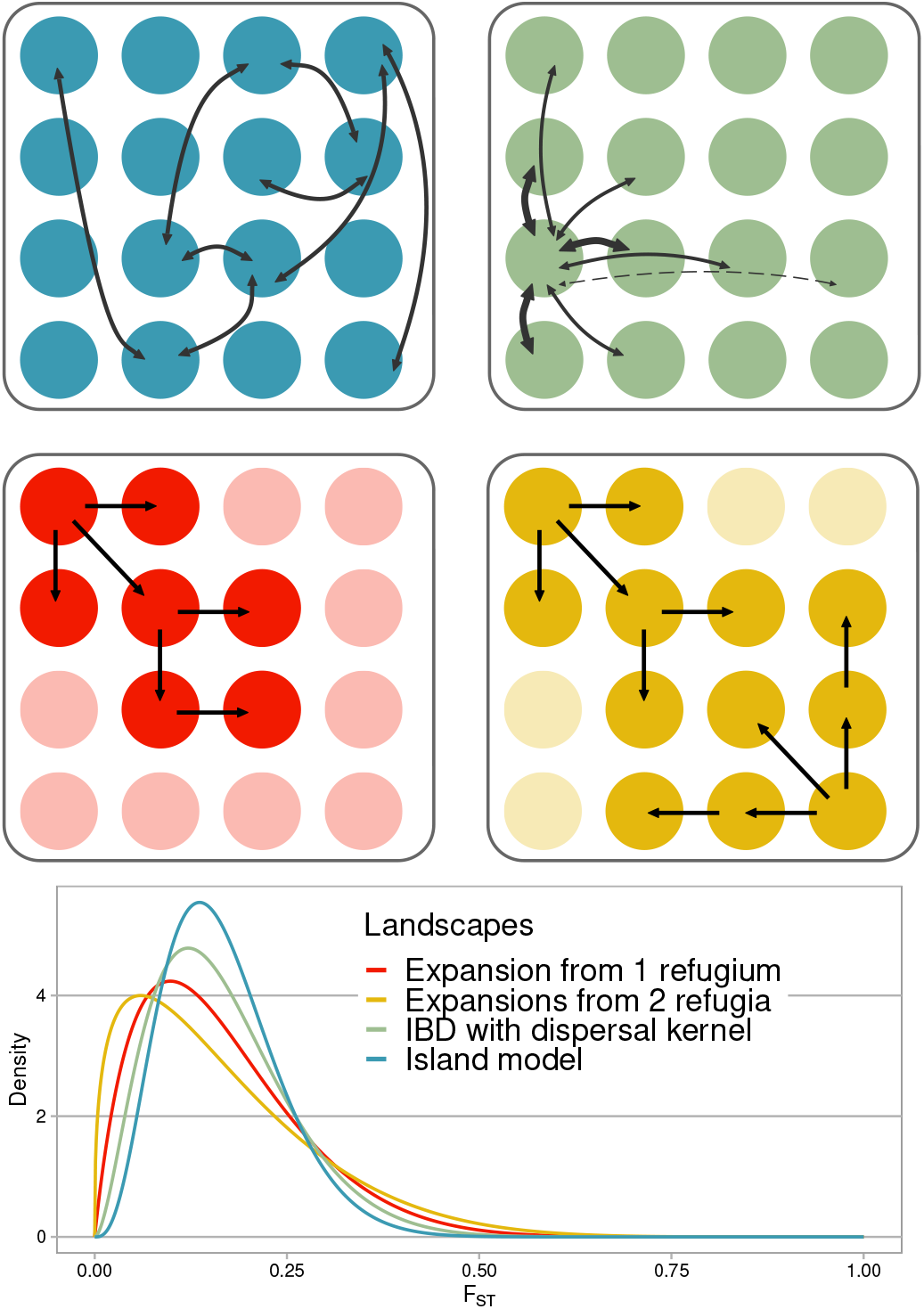
Importance of demographic history on the null genomic distribution of *F*_ST_. Represented here are four metapopulations of *d* = 16 subpopulations/demes (filled circles) differing in their demographic history (adapted from [12]. Blue: island model where each deme is at carrying capacity, and equally connected by gene flow to all other *d* - 1 demes (arrow). Green: isolation by distance model with two-dimensional dispersal kernel. Red: range expansion model from a single refugium (top left deme). Arrows indicate here the colonisation of previously free niches. Yellow: range expansion model from two refugia (top left and bottom right). Curves represent the genomic distribution of *F*_ST_ under each model. The four models produce different genomic *F*_ST_ variation for the same mean. In particular, areas beyond the blue curve represent false positive *F*_ST_ outlier regions if the island model is taken as the null model.

Hence, the variance of *F*_ST_ across genomic regions can be strongly increased according to the demographic scenario experienced by the studied organism. For instance, for the same genomic average of *F*_ST_, a two-dimensional isolation-by-distance model (with a dispersal kernel) will increase the genomic variation when compared to the island model where all sub-populations are equally connected by gene flow. Additionally, this genomic variation can further increase as a result of colonization of empty niches from one or multiple refugia (Fig. 1). These expansions are important processes in the history of species in response to continuous cycles of historical climate fluctuations, impacting both marine and terrestrial organisms [13]. Recent polyploid species are frequently more present in invasive ranges than their closest diploid relatives [14, 15], suggesting that the island model is even less likely to apply to them. Neglecting such demographic processes will therefore lead to an increase in the false discovery rate when searching for genomic regions involved in local adaptation, if the classic island model is blindly applied to a studied organism without evaluating its relevance. Riverine or coastal organisms are thus not expected to share the same null envelope as pelagic organisms for the same genomic average *F*_ST_ [16]. Taking the specific demographic history of the studied organism into account is then crucial to reduce the number of genomic regions erroneously claimed to be involved in local adaptation. Fortunately, methodological developments in population genomics since the early 2000s have greatly facilitated the exploration of sequenced data to capture the main demographic features behind the observed genetic patterns. Because nowadays the whole ABC analysis can be done in four short command lines in a Linux terminal, then the aim of this chapter is not to detail one particular inferential tool among the many available to date (for more details on the current state of the art, see the review by [17], but to detail the steps that lead to the exploitation of the great flexibility of ABC (approximate Bayesian computation; [18]) in particular to tackle the difficult issue of exploiting polyploid genomic data, and by extension, all approaches based on comparisons between simulations and observations.

## 2 Methods

### 2.1 Demography inferred from population genomic data

It is important to begin this chapter by defining what demographic inference is, particularly to avoid confusion with phylogenetic reconstruction. Firstly, the time scale involved is not the same: phylogenetic reconstruction aims to group divergent organisms according to their relative divergence for certain molecular and/or morphological metrics, and then to propose a temporal sequence leading to the established groups. In contrast, demographic reconstruction aims to describe the temporal changes in patterns of migration and in effective population sizes that occurred within a system of closely related species. In the early days of molecular population genetics, relationships between genetic patterns and demographic processes were proposed. For instance, since the per generation probability of having a coalescent event in a sample of gene copies is proportional to 1/2Ne, where *Ne* is the number of diploid individuals making the population, then the total size of the sample pedigree increases linearly with *Ne*. By assuming a mutation model with an infinite number of sites, it follows that the number of SNPs in a population, for a given neutral locus, increases linearly with the size of the coalescent tree, and hence, with *Ne*. Thus, assuming a population whose size was constant in its recent history, it became possible to estimate this value of *Ne* from the number of segregating polymorphic sites observed at the sequenced neutral loci [19]. In addition to this relationship with the number of SNPs at a given neutral locus, *Ne* is also linearly related to the expected average number of nucleotide differences between pairs of sequences per site (*π*; [20, 21]. The effective number of individuals in the population therefore directly impacts both the number of SNPs observed in a given nucleotide sequence alignment, but also the distribution of allelic frequencies. Deviations from this standard demographic model of constant population size leave different signatures in the genomes, including the effects of variations in population sizes over time [22, 23], migration between populations and/or introgression between species [24, 25] and ancient population structure [26]. Demographic inferences thus aim to control for the influences of different processes on patterns of intra-population polymorphism and on patterns of interspecific divergence by capturing the signatures they have left in the genomes. By integrating these different genomic signatures let by *Ne* variation, genetic structure and introgression into an inferential framework, it becomes possible to interpret sequence data in the light of models deviating from the standard neutral model (Fig. 2).

**Fig. 2.**
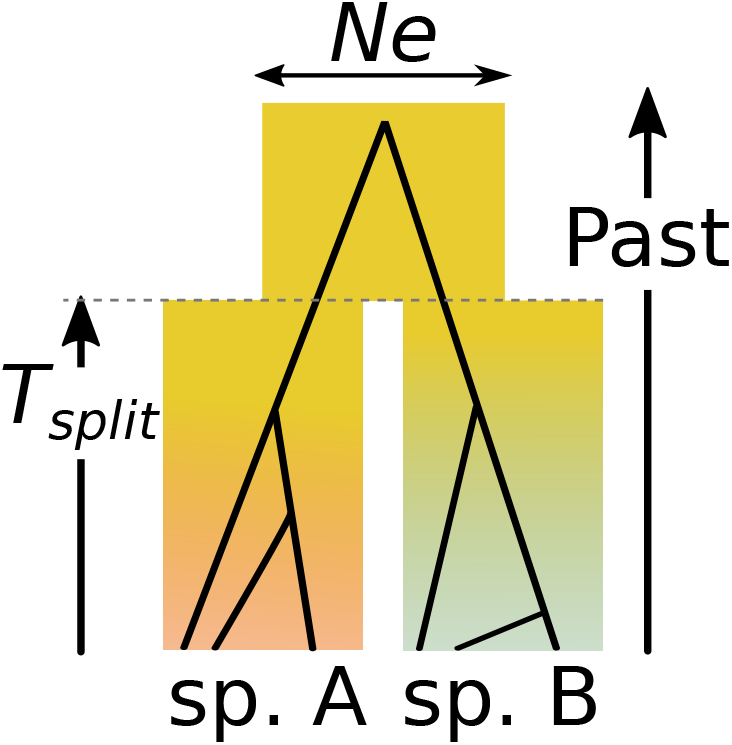
Demographic model of divergence from a common ancestor. In its simplest version, the model describes the subdivision, *T*_split_ generations ago, of an ancestral population with *Ne* individuals into two closely related populations giving the future species A and B. This demographic model can be extended to add i) population structure and ii) migration between different populations at different times.

These more complex demographic models can be defined by sets of parameters that constrain the evolution of one or more populations/species. For instance, four parameters are sufficient to describe the simplest model of divergence of two gene pools: the divergence time *T*_split_ (in number of generations) and the effective sizes of the three populations (the ancestral one plus its two daughters). Under this model, the age of subdivision of the ancestral population will affect the genomic *F*_ST_ between the two diverging daughter populations A and B in different ways. Firstly, *F*_ST_ increases monotonously with divergence from zero to one in the absence of continuous migration or secondary contact between the diverging daughter populations (Fig. 3-A). Secondly, the variation in *F*_ST_ across loci, initially low in the early stages of divergence, will increase under the effect of drift and recombination, generating genomic heterogeneity in differentiation. Stochastic and progressive fixation of different alleles between the two populations at different positions (initially from standing variation, then, from derived mutations) finally tends to reduce the variation of the *F*_ST_, producing somewhat homogeneously differentiated genomes (Fig. 3-B).

**Fig. 3.**
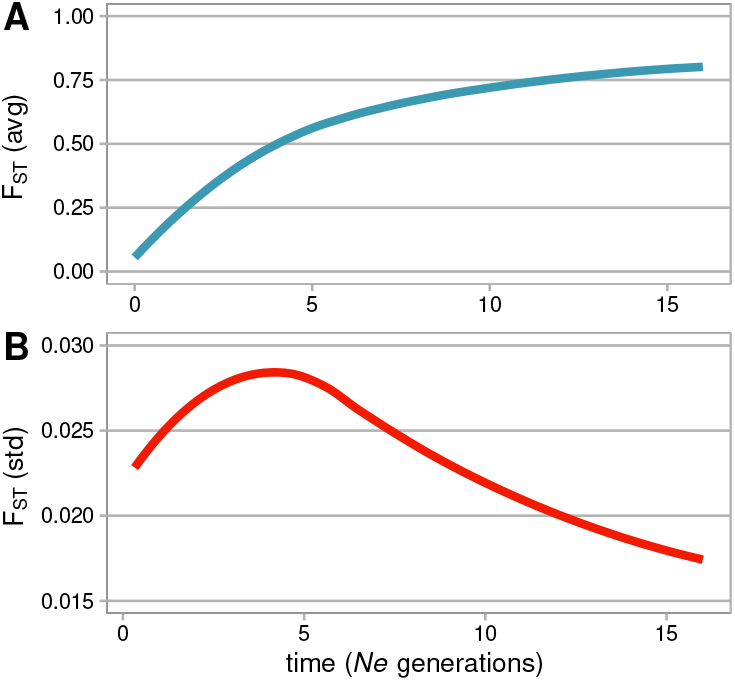
Evolution of *F*_ST_ during divergence. **A.** monotonic increase in the average genomic *F*_ST_ over time, expressed here in *Ne* generations. **B.** Genomic variation of *F*_ST_ initially increases from genomically homogeneous populations at low levels of differentiation and then decreases to genomewide differentiation.

The patterns of diversity encapsulated in the two-dimensional Site Frequency Spectrum (SFS) also evolve as a result of genetic drift over time (Fig. 4). In the first generations immediately after the split a strong correlation in allele frequencies is expected between closely related populations (Fig. 4 for *T*_split_ = 0.0001*Ne* generations). At this level of divergence, the majority of SNPs correspond to shared polymorphic sites between the two populations, inherited from the ancestral population. Due to the sampling effect, some SNPs may appear to have fixed an allele in one population that still segregates at high frequency in the other population. During the divergence process, the proportion of genotyped SNPs exhibiting shared polymorphisms in intermediate frequencies decreases (*T*_split_ = 0.4*Ne* generations), until it almost disappears (for *T*_split_ > 1*Ne*) to the relative benefit of polymorphisms specific to one of the two populations, and finally, to fixed differences between them [27].

**Fig. 4.**
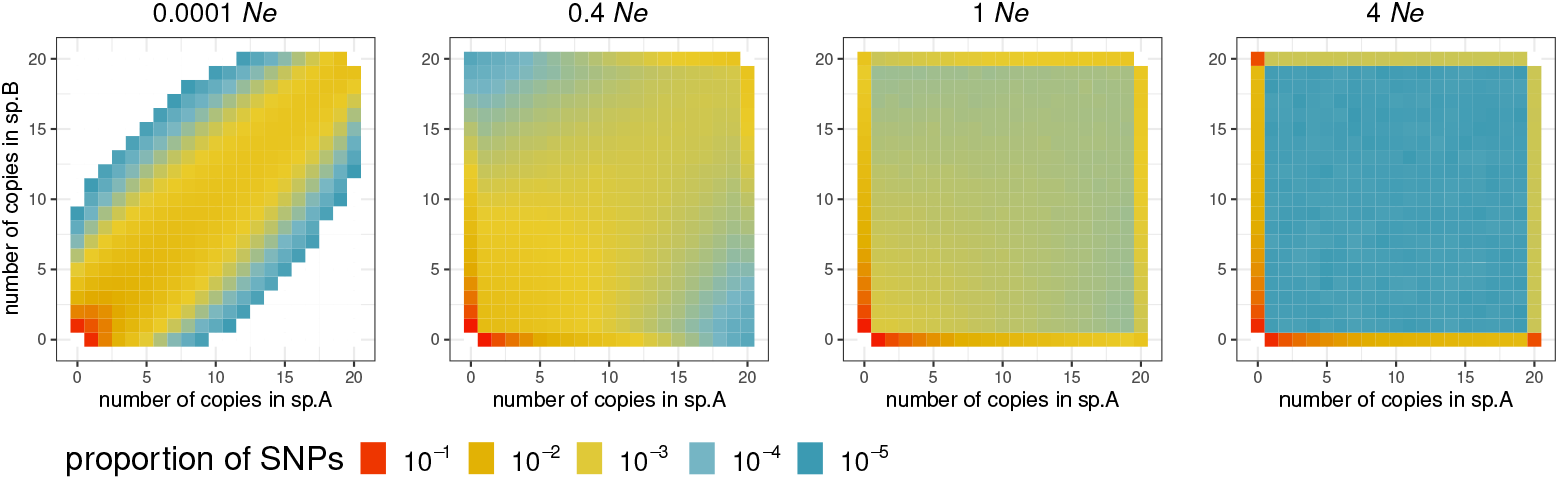
Joint allelic frequency spectra (jSFS) during divergence. Four spectra describe here the allelic frequencies in two diploid sister species at four different divergence times expressed in *Ne* generations, where *Ne* is the effective population size of the two daughter populations and their common ancestor. From left to right: *T*_split_ = 0.0001*Ne* generations ago, 0.4*Ne*, 1*Ne* and 4*Ne*. These spectra are formed from 20 haploid gametes sampled and sequenced from the two diploid populations/species A and B. For each SNP identified among the 40 sequences, the number of copies of the derived allele found in species A (x-axis) and B (y-axis) is counted. Colours indicate the expected proportions of genotyped SNPs falling within each bin of the jSFS. Shared polymorphisms are located in the inner square. Polymorphisms that are private to a single species are in the external contour. Derived mutations that have become fixed in a single species are in the lower right or higher left corners.

Exploiting information from this two dimensional SFS was first proposed by Li and Stephan [28] in order to reconstruct the demographic history of Drosophila melanogaster. They showed that the jSFS contained the information necessary to capture major historical events: *i*) past population expansions in Africa; *ii*) relatively recent establishment of *D. Melanogaster* in Europe; *iii*) a strong bottleneck associated with this colonisation. Other methods have since followed up on this approach by fitting demographic parameters to the jSFS, either by maximum likelihood [29, 30, 31] or by Bayesian approximation [32] procedures. More recently, the information contained in the SFS has allowed alternative models of speciation to be compared in order to highlight a secondary contact between the Mediterranean and Atlantic lineages of the sea bass [33]. While these methods of reconstructing demographic histories from nucleotide sequencing data have since been widely applied to diploid organisms, some work has shown that they can also be successfully extended to tetraploid species [34, 35]. The following section will therefore detail how to apply such methods to tetraploid organisms, and how to take advantage of their methodological flexibility to adapt to certain biological properties.

### 2.2 from tetraploid sequences to first inferences

To date, demographic inferences using ABC approaches have largely been applied on diploid species, notably to: *i*) test demographic expansions and/or contractions [36]; *ii*) test whether two populations/species are connected by ongoing gene flow [37]); *iii*) test different demo-genetic models, e.g. to take into account the effects of indirect selection on the genomic distribution of diversity [38, 39], or to identify molecular targets of selection against hybrid incompatibilities [40, 41]. In addition to these issues which are relevant for both diploid and polyploid species, other questions specific to polyploids arise. One is often interested in distinguishing between the establishment of a polyploid lineage following the doubling of its genome within a single population (autopolyploidization) and its establishment as a result of hybridization associated with genome doubling between two divergent populations/species (allopolyploidization). Although the distinction between auto- and allopolyploidy is conceptually important to gain insight into the origin of a given tetraploid species under study, it is often achieved indirectly by deducting the evolution mechanism from the mode of inheritance. Usually, disomic inheritance, i.e, preferential pairings between only two homologous chromosomes, is interpreted as indirect evidence of an allopolyploid origin. In contrast, polysomic inheritance (random pairings among all homologous chromosomes) is interpreted as indirect evidence of an autopolyploid origin. But tetrasomic inheritance in a recently duplicated tetraploid lineage will generally gradually evolve to disomic inheritance [42], a process called “diploidisation”. This process results from the progressive divergence between pairs of homeologous chromosomes, accelerated by reduced local recombination rates (i.e, in case of large chromosomes), as well as by the subfonctionalisation and neofonctionalisation processes acting on homeologous genes. Because this process of diploidisation can be prevented at islands of recombination between the two sub-genomes [43], differences in gene exchange between homeologous pairs across the genome can lead to mixed or intermediate inheritance models that are called “heterosomy”, with some loci showing disomic and others tetrasomic inheritance. Such mixed inheritance models are somewhat neglected in the population genomic literature despite being well reported [44, 45, 46]. The evolution of disomy in autotetraploids suggests that inferring the mode of polyploidisation from the mode of inheritance only may be misleading, but there is much to learn by decoupling these properties. Here we describe how this can be achieved by detailing key steps in demographic inference in tetraploids (Fig. 5):

1. Designing the models to be explored.
2. Processing of observed and simulated sequences.
3. Comparison between observation and simulations.
4. Checking the results.

**Fig. 5.**
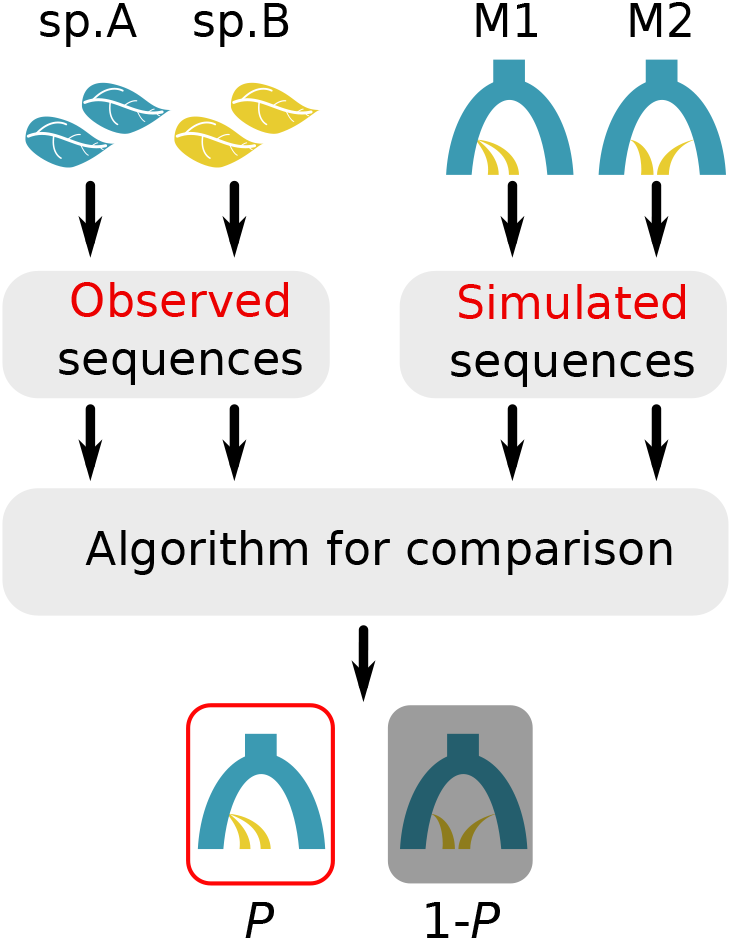
Three steps in ABC for demographic inferences. The observed data is obtained by DNA/RNA sequencing of individuals from natural populations/species A and B. Sequences are then summarised by a vector of chosen statistics to quantify the genomic patterns of polymorphism and divergence. Simulated data are exploited to obtain null joint-distributions of the same summary statistics under each of the demographic scenarios to be evaluated. The last step in model classification is a statistical comparison between simulated and observed statistics in order to assign a probability *P* to the best supported model. *P* quantifies the relative abilities of the compared models to produce simulations close to the observation.

#### 2.2.1 Designing the models to be explored

The principle of ABC-like approaches is to first classify an arbitrary set of models according to their ability to reproduce the observed data. This requires upstream design of the alternative scenarios that an experimenter wishes to evaluate. It is therefore crucial to put the hypothesis to be tested at the centre of the design. In the speciation genomics literature, one of many questions of interest is whether or not there is ongoing gene flow between two divergent species. So the experimenter proceeds to design alternative demographic scenarios that may or may not lead to species currently connected by gene flow [47, 48, 49, 50]. A first question of interest concerning polyploids could, for instance, be the auto- *versus* allopolyploid origins (Fig. 6). In such models, each sub-genome making a tetraploid taxon is considered to be a diploid population. Autopolyploidisation therefore involves simulating the divergence of a diploid species into three diploid lineages at a time *T*_WGD_ [34]: one lineage resulting in a daughter diploid species (species A in fig. 6-A), which is closely related to the other two lineages making the daughter tetraploid species (species B in fig. 6-A). Similarly, allopolyploidisation is simulated by modelling two divergence events taking place simultaneously at time *T*_WGD_, one between parental species A and sub-genome A of species B, and, the other between parental species C and sub-genome C of the tetraploid species B (fig. 6-B).

**Fig. 6.**
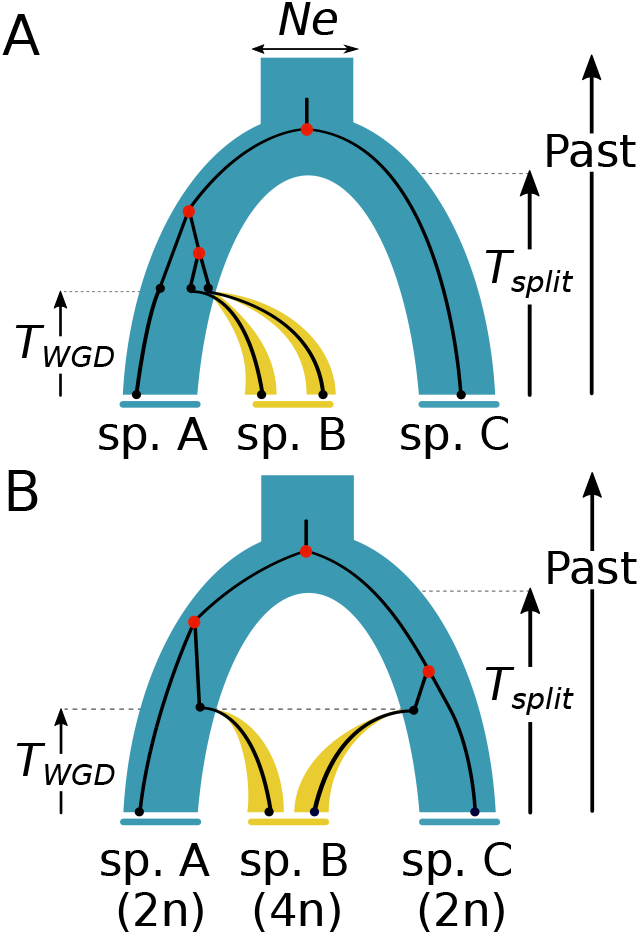
Demographic speciation scenarios with polyploidization. Blue: diploid species/populations. Yellow: tetraploid species/populations, resulting from **A** autopolyploidization or **B** allopolyploidization occurring *T*_WGD_ generations ago. Each of the two sub-genomes forming the tetraploid taxon are simulated as two diploid populations with or without gene flow among them to mimic tetrasomic *versus* disomic inheritance, respectively. A strict bottleneck is assumed here at the origin of the tetraploid with a single founding individual, followed by exponential population growth.

This is an opportunity to raise a central point in model design: a demographic hypothesis test frequently involves a test between at least two different categories of scenarios that differ for the hypothesis to evaluate. In the popular example of a test for recent introgression between two species of interest, several scenarios can lead to ongoing gene flow. Not exclusively, ongoing gene flow between two species can be the result of secondary contact between them after a past allopatric divergence. But such ongoing gene flow may also have occurred continuously between these two lineages, without interruption, since their divergence from their common ancestor. Similarly, the absence of migration between two species may be the result of a reduction in gene flow which may be sudden or progressive over time. Hence, testing the hypothesis of current gene flow between two species would consist in comparing two categories of scenarios: secondary contact or continuous migration *versus* sudden interruption or progressive interruption of gene flow. Categorising the hypotheses to be tested in this way allows possible confounding effects to be taken into account, which, when neglected, can bias the biological conclusions drawn from the inferences [39]. Applied to the elucidation of the mode of polyploidisation, several sub-models can compose each of the two categories auto- and allopolyploidisation. Both categories can be simulated assuming tetrasomic autopolyploids only and disomic allopolyploids only. In this case, an ABC approach will risk interpreting a functionally disomic autopolyploid as a species with allopolyploid origin. Assessing the mode of polyploidisation therefore requires implementing the different modes of inheritance (disomic, heterosomic, tetrasomic) for both origins [35]. This is an illustration of the power of simulation-based approaches such as ABC, where confounding effects (for instance: disomic = allopolyploid) can be avoided by regrouping variants of the same scenario. This is now possible thanks to the great flexibility offeredby the various simulators available [51, 52, 53, 54, 55, 56, 57, 58, 59]. Because the two diploid sub-genomes constituting the tetraploid species can be simulated as two diploid populations, then, the process of recombination between the subgenomes under tetrasomic inheritance is equivalent to infinite gene flow between sub-populations. In contrast, disomic inheritance is equivalent to a strict isolation between sub-populations. In between, mixed inheritance (heterosomy) corresponds to a proportion *α* of the genome where homeologous chromosomes are connected by gene flow, and 1-*α* are isolated. Hence, if *α* is assumed to be equal to 1 then inheritance is strictly tetrasomic, and strictly disomic if one assumes *α*=0. Detecting whether a species is autopolyploid or allopolyploid using ABC or supervised deep learning approaches requires, at a minimum, simulating 6 different demographic scenarios, corresponding to all combinations between origin ([autopolyploidy, allopolyploidy]) and inheritance ([disomy, heterosomy, tetrasomy]) [35]. Recently, Booker et. al. [60] have gone further by adding different migration sub-scenarios within these two categories and successfully applied this approach to hundreds of loci obtained using exome capture from the North American gray treefrogs Hyla. They test the same 6 models, with the addition of: 1) models with gene flow from *Hyla chrysoscelis* (diploid) to *H. versicolor* (tetraploid); 2) gene flow in both directions (but at non-necessarily symmetrical rates). Using an ABC approach, they strongly reject models with strict reproductive isolation between these two frog species suggesting introgression is effective despite differences in ploidy levels. It is important to be mindful that while the ABC or deep learning method will provide greater support to one of the proposed models (or categories of models) relative to the others, this relative support depends solely on the arbitrarily chosen set of alternative models. By analogy, the ABC will strongly support the “12-sided die” model in a 12 *versus* 6-sided comparison, if the actual model is the untested 20-sided die. The philosophy of this approach, in essence, is to classify models relatively to each other. Thus, the interpretation of the results must always be expressed in the light of the set of compared models rather than as definitive conclusions decoupled from the initial model design. In our initial study [35], we proposed a set of models for which a sample mimicking that of the experimenter (in terms of number of individuals, number and length of sequenced loci) would be simulated for a number of replicates set by the user. The purpose of such simulations is to empirically produce joint distributions of *N* summary statistics (more details about them in the next section) under the different scenarios to be evaluated. In practice, this is done by running the script *run_ABC_polyploid_v2.py* of our *ABC_WGD* pipeline (https://github.com/popgenomics/ABC_WGD). Our intention here being to accompany the new user in the logic of the analysis, but we invite the interested reader to follow the Readme of the GitHub repository for a more technical tutorial that will be adapted to possible future updates.

#### 2.2.2 Processing of observed and simulated sequences

ABC methods are based on the comparison between simulations performed according to different alternative models and observations derived from the analysis of real data. The comparison is made through summary statistics which are used to describe the observed patterns of polymorphism and divergence genome-wide: the model receiving the strongest relative support being the one that produces combinations of summary statistics closest to the observed data. A step that becomes crucial is therefore the choice of relevant summary statistics. We will not develop here the issue concerning the number of statistics to choose, as major improvements brought by Random Forest algorithms now allow the use of a large number of them without loss of power [61]. Nowadays, the experimenter wishing to apply a custom ABC approach needs to focus on two important aspects of the choice of summary statistics:

1. Which statistics best capture past demographic events?
2. Which statistics can be quantified on the observed dataset (i.e. statistics requiring phased data? Is the identification of parental sub-genomes required? etc…)

A strength of ABC is its adaptability to different sampling and sequencing strategies. This flexibility circumvents problems that arise when calculating summary statistics for a polyploid. One of them is to phase the haplotypes, and to attribute each sequenced copy to one of the two parental sub-genomes. We proposed in 2015 [35] a simple approach, with the aim of being transferable to the widest range of sampling and sequencing strategies (and ploidy levels), by using only summary statistics based on allelic frequencies (figures 7 and 8).

**Fig. 7.**
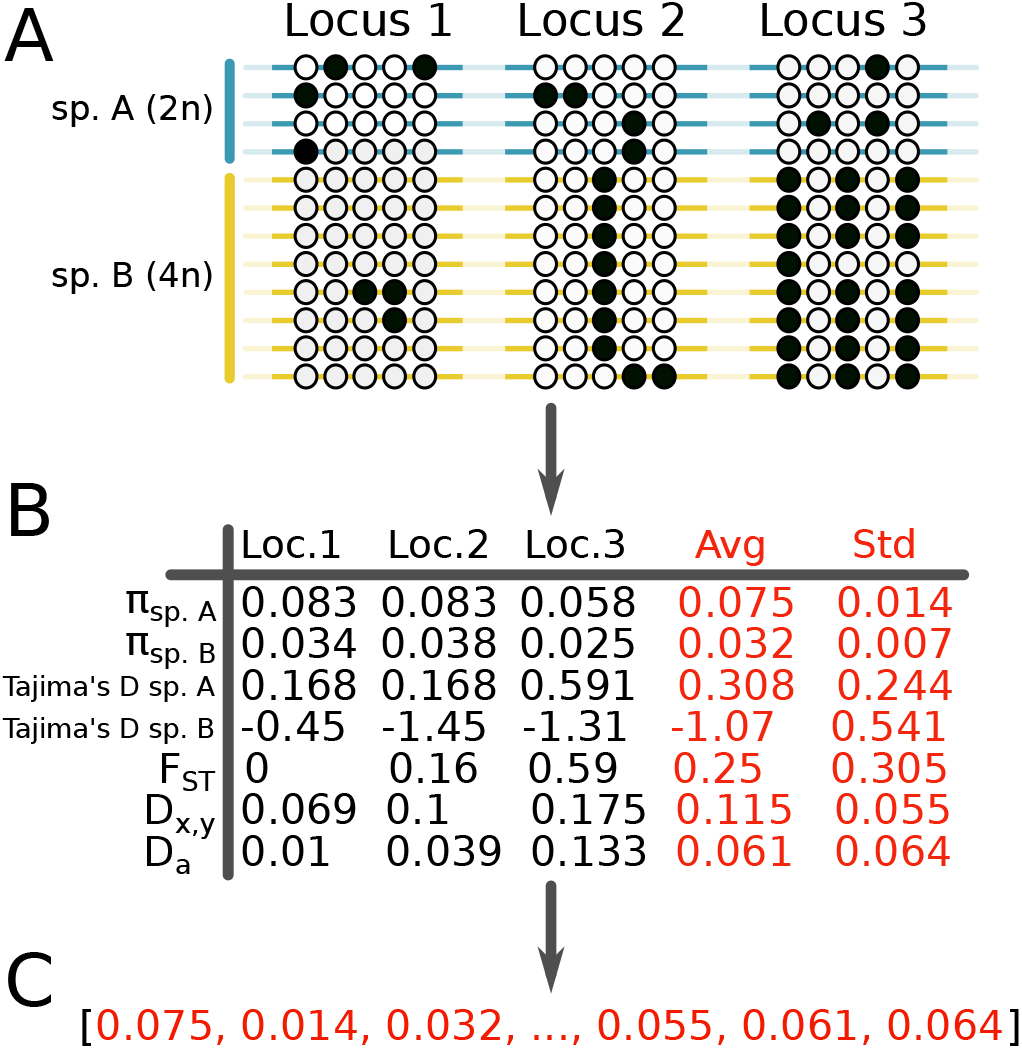
Summarizing the data by using classical statistics in population genetics. Three loci were sequenced in two diploid (species A in blue) and two tetraploids (species B in yellow) individuals. The white and black circles indicate alternative alleles at the identified SNPs. A set of 7 summary statistics is calculated for each locus. The vector containing the means and standard deviations of each statistic will be used as the input for inferences.

**Fig. 8.**
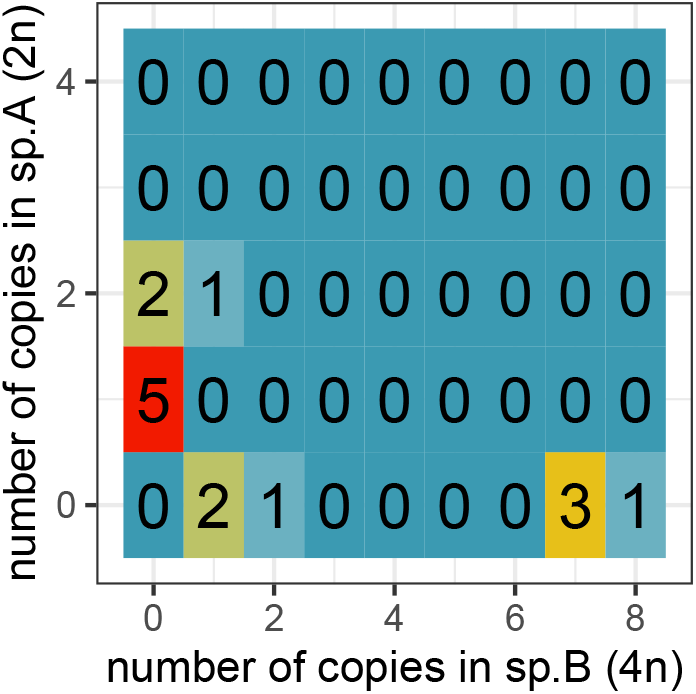
Joint site frequency spectrum as summary statistics. The example found in figure 7-A is now summarised by an SFS, where each bin corresponds to the allelic count of the derived allele (black) within the sampled and sequenced populations/species. The number of SNPs within each bin can be used as a statistic for inferences.

This approach assumes having a good confidence in the allelic count for each SNP, regardless of the ploidy level. This means an absence of sequencing bias, where all of the four copies of a gene carried by an auto- or allotetraploid individual are taking into account when assessing the genotype. Different tools propose to genotype sequencing data from polyploid organisms, by dealing more or less efficiently with sequencing errors, allelic bias and overdispersion, among them: the method of Li [62, 63], GATK [64], fitPoly [65] and updog [66]. Once the sampled genotypes are genotyped, it becomes possible without phasing to define a minimum but still efficient set of statistics suitable for demographic inference, among them: *π*; Watterson’s *θ*, Tajima’s *D*, *F*_ST_, raw divergence *D_x,y_*, net divergence *D_a_*. The purpose of these statistics is to quantify the patterns of polymorphism and divergence, in order to find the model that best reproduces them. Experimenters wishing to apply/develop an ABC approach to studying the history of their model organism must bear in mind that there is no single ‘magic’ statistic that will resolve on its own the history of an organism: demographic signatures are essentially retrieved in the joint combination of different statistics. More precisely, demography impacts the genomic distributions of different statistics [67]. Quantiles of these genomic distributions, usually mean and standard deviation, should then be used as summary statistics, although any method describing a distribution would be relevant (figures 7-A and 7-B). A dataset consisting of several sequenced loci in different individuals from related species is finally summarised into a vector of statistics (Fig. 7-C) from which inferences will be made, in order to propose the simulated model-parameter combination reproducing the best this vector of observations. In addition to standard population genetics statistics, SFS can also be treated as a vector of summary statistics (Fig. 8). The latter does not require phased data, nor attribution to one of the parental subgenomes, making it applicable to genomic data in tetraploids. Each bin making the SFS becomes an independent statistic summarising the observed data.

The representation of closely related diploid and tetraploid populations in the form of a SFS highlights the effects of modes of inheritance on genomic patterns (Fig. 9). During autopolyploidisation with tetrasomic inheritance, the bottleneck associated with the emergence of the tetraploid taxon rapidly purges a large amount of the polymorphism segregating in the ancestral population. As a result of such strong bottleneck, each bin of the inner square (i.e, corresponding to shared polymorphism) or on the bottom edge (i.e, polymorphisms exclusive to the tetraploid lineage) of the SFS represents about a proportion of 0.0001 of the total number of SNPs found in a diploid/tetraploid alignment 0.01*Ne* generations after genome duplication. In this autopolyploidisation scenario, the first tetraploid individual is the result of a panmictic mating within the parental diploid population. The associated bottleneck thus generates a peak of heterozygosity (representing about twenty per cent of the genotyped SNPs) in the generations immediately after genome duplication, but which will quickly fade as a result of recombination associated to unbiased Mendelian segregation. At this same stage in the divergence process, the most represented category of SNP is the polymorphism segregating only in the diploid species (Fig. 9-A, left edge). Although the polymorphism within the newly established tetraploid population represents a very small part of the genome, alleles can segregate in frequencies between zero and one due to the recombination between the four copies. From the early stages of divergence, genomic patterns differ between tetrasomic autopolyploid and disomic allopolyploid. In the latter, the most prevalent category of SNPs is the stabilisation of alleles at frequency 0.5 in the tetraploid (Fig. 9-B).

**Fig. 9.**
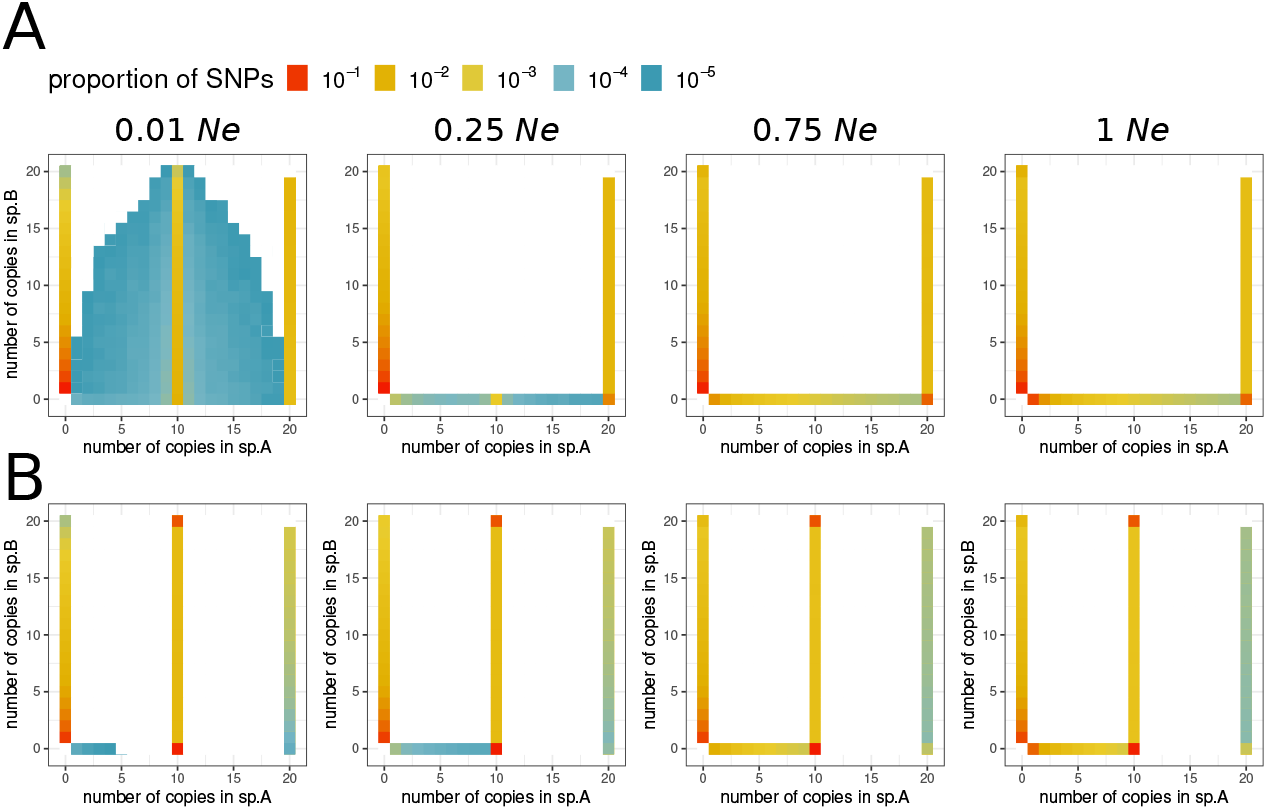
Joint allelic frequency spectra (jSFS) after whole genome duplication. Joint spectra describing the distribution of allelic frequencies for a pair of diploid (10 sampled individuals) and tetraploid species (5 sampled individuals). The genome duplication took place *Ne* generations after the separation of the ancestral population. The four jSFS represent genomic patterns after 0.01, 0.25, 0.75 and 1 *Ne* generation after the **A**. autopolyploidization (tetrasomic) or **B.** allopolyploidization (disomic).

The other difference is the absence of alleles segregating at frequencies above 0.5 in the disomic allotetraploid. Hence, the process of polyploidisation has a drastic immediate effect on polymorphism patterns. The time since genomic duplication will mainly contribute to increase the relative importance of segregating polymorphism in the tetraploid. These marked differences in the SFS mean that an ABC approach will be able to correctly distinguish a tetrasomic autotetraploid from a disomic allotetraploid with near 100% accuracy. If all three types of inheritance (disomy, heterosomy, tetrasomy) are taken into account for each of the two origins (auto and allo), then the rate of identification of the correct auto- *versus* alloploidization remains high, but is strongly impacted by the extent of genomic tetrasomy. Thus, ABC will correctly support the (auto or allo) tetraploid origin in ~ 99% of cases if the species is disomic, in ~ 90% of cases if the species is heterosomic, and ~ 80% of cases [35] if the species is tetrasomic. Increasing the extent of recombination between homeologous chromosomes leads to a loss of information about the mode of polyploidization. This is a consequence of genetic drift, which increases the possibility of losing one of the two parental genomes when the new tetraploid population is still restricted to a few individuals. This biological explanation of the error rate of the ABC in classifying models puts it into perspective. Indeed, if one of the two parental genomes has been lost by drift in the early generations of a sufficiently tetrasomic allotetraploid population, then the species is genomically autotetraploid.

#### 2.2.3 Comparison between observation and simulations

In order to make statistical inferences using an ABC approach, three objects are needed:

1. A vector of summary statistics obtained from the observations (Fig. 7-C).
2. A reference table obtained from the simulations, with columns corresponding to the vector of summary statistics and rows corresponding to different simulation replicates.
3. A vector of the variable to be estimated (categorical indicator variable in the case of a model comparison; continuous variable in the case of a demographic parameter under a given model), in correspondence with rows of the reference table.

The reference table is obtained thanks to the pipeline described in the previous sections in order to simulate the experimenter’s sample (number of individuals, number of sequenced loci/chromosomes) under the demographic scenarios to be studied and, to obtain summary statistics from the performed simulations. While the aim of this book chapter is to encourage experimenters to appropriate existing simulation tools in order to develop a pipeline for their biological questions, such a project may scare neophytes. For this reason, we have developed a turnkey version of such a pipeline, freely available on GitHub: https://github.com/popgenomics/ABC_WGD

The *run_ABC_polyploid_v2.py* command alone will produce the reference table needed for downstream analysis by combining two output files named *ABCstat.txt* and *ABCjsfs.txt*, for different combinations of polyploidisation and inheritance modes. A third file is produced, *priorfile.txt*, which contains the values of parameters, randomly drawn from appropriate distributions according to the priors set by the experimenter, which were used to produce each simulation replicate. The command *run_ABC_polyploid_v2.py* simply takes as arguments the name of the model to be simulated (auto, allo), the inheritance mode (disomic, heterosomic, tetrasomic), the limits of the parameter values to be explored (priors) and the number of multilocus iterations. The reference table (obtained by combining *ABCstat.txt* and *ABCjsfs.txt*, if *run_ABC-polyploid-v2.py* from our pipeline is used) is composed in columns by the same summary statistics than the statistics used to describe the observed dataset (vector of summary statistics), one row per simulated iteration. There are no magic numbers concerning the summary statistics or the number of iterations per scenario to be tested. In practice, it is usual to see in the literature between 10,000 and 50,000 iterations per scenario if a random forest algorithm is used downstream (e.g. R library *abcrf* [68], and around 1 million iterations if a rejection/regression algorithm is preferred (e.g. R library *abc* [69]. It is up to the experimenter to explore around these orders of magnitude to test whether the number of iterations has an effect on the accuracy of *M*_0_ (Model with autopolyploidization) *versus M*_1_ (Model with allopolyploidization) comparison. Regarding the number of summary statistics, typically less than 50 are used, but this number can be greatly increased by including each SFS bin in the list of statistics to be used, which is a function of the sampling size. In the case of a model comparison, the third required object is a vector containing the model indicator variable. This vector, of the same length as the number of rows in the reference table (containing the statistics simulated under all models compared, such as *M*_0_ and *M*_1_), is intended to label each row in the reference table with the actual model (e.g., either *M*_0_ or *M*_1_) used to produce the row in the table. Using the *abcrf* package in R, once these three objects are produced, the comparison of *M*_0_ *versus M*_1_ models is done in two simple and rapid steps, corresponding to two short R functions to execute. The first step is a training step of the random forest (using the *abcrf()* R function, from the eponymous package), involving the reference table and the vector with the model indicator variable. The returned object is a random forest of *ntree* decision trees, trained to predict the model that best explains a given vector of summary statistics. The effect of the *ntree* value on the accuracy of the *M*_0_ *versus M*_1_ comparison can be empirically explored. To provide an indication, a value of *ntree* = 1, 000 appears in the literature as standard but may vary depending on the number of models compared and the number of summary statistics. The second step is a prediction step (using the *predict()* R base function) that uses the random forest trained in the previous step and the vector of observations. This step returns three values of interest:

1. The selected model (*M*_0_ or *M*_1_ in the case of a comparison between two models; but *M*_0_ and *M*_1_ can also be categories of models).
2. The number of trees in the forest that voted for each of the models compared.
3. The posterior probability of the best model (or the best category of models), approximated by 1 - *ϵ*, where *ϵ* is the error rate of the inference.

Once the best scenario among those proposed has been identified using the *abcrf()* and *predict()* functions, the next step is to estimate the parameter values that best explains the observed data (of the selected model). Again, using the *abcrf()* R library, this can be achieved in two short steps. The major difference with the previous step of model comparisons is that the objective here is to predict a continuous variable (the model parameter to be estimated) and not a categorical variable (the model *M*_0_ or *M*_1_ to be inferred). To do this, a random forest must first be trained using the same reference table as previously for the model comparison, but replacing the model indicator vector with a vector containing the value of the parameter X that one is attempting to infer. The function *regAbcrf()* from the R package *abcrf* will build a regression random forest from the simulated statistics and the used parameters. For the second step, the *predict()* function will, taking as objects the trained regression forest and the vector of observed statistics, predict the expected value of the parameter, the variance of the posterior and its quantiles. To provide an idea about the computation time required, we have here estimated the time of whole genome duplication (*T*_WGD_) for the four combinations “autopolyploidization or allopolyploidization” x “disomic or tetrasomic”. For each of these four combinations, we simulated 10,000 datasets to train the regression forest, and 30,000 datasets to test the trained forest (Fig. 10). On a 2017 laptop, with 6 dedicated threads, learning the four forests and estimating *T*_WGD_ for the 4×30,000 pseudo-observed datasets [70] took 13 minutes. This execution time is not impacted by the sample size in terms of number of individuals and loci. Such a re-analysis of simulated data sets (not used this time to train the forest) is necessary to assess the quality of inference along the range of explored values. In this way, we show that the age of polyploidization *T*_WGD_ can be accurately estimated for recent, intermediate and ancient events, and for each of the four explored models (Fig. 10).

**Fig. 10.**
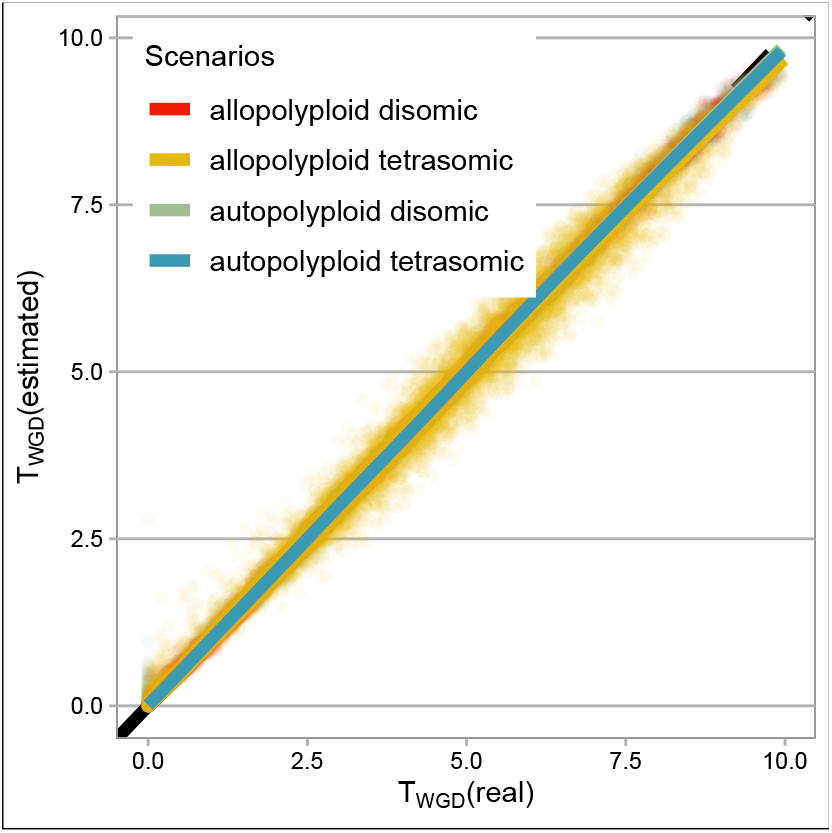
Test of accuracy in estimating *T*_WGD_ for four models. It is assumed here that the observed dataset consists of 5,000 loci of 1 kb, sequenced from 20 diploid individuals and 10 individuals from another tetraploid species. The age of polyploidisation, (expressed in *Ne* generations), as well as all the parameters of the model, is randomly drawn between 0 and 10. The objective of the accuracy test is to use a first set of 10,000 simulations to train the regression forest, and a second set of 30,000 simulations to empirically establish a relationship between the real value of *T*_WGD_ (x-axis) and the value predicted by the trained forest (y-axis). Points represent the 120,000 estimated values for *T*_WGD_. Lines represent the loess regression between real and estimated values. Black line has a slope of 1 and an intercept at (0, 0). Colors represent the four explored models.

## 3 Conclusion

While turnkey pipelines are nowadays available in order to efficiently infer the speciation history of tetraploids from genomic data (https://github.com/popgenomics/ABC_WGD), it is still important to highlight that the strength of ABC-like approaches is their flexibility. Thus, pre-existing pipelines can be immediately useful for analysing newly acquired data, but more importantly, they can be useful as a basis for customisation. A first layer of customisation may involve the demographic models to explore, either by adding genomic introgression, or by changing the population growth function of the neo-polyploid, or by increasing the ploidy level, etc… A second layer of customisation can be applied to the genomic models, notably by implementing more relevant inheritance patterns, different natural selection models. A third layer of customisation may concern the set of summary statistics, notably by implementing those that can be brought to light in future. And finally, the prediction algorithm itself can be modified, notably by taking advantage of the promise of convolutional neural networks in population genomics [71]. As with every inferential method, it is always important that the experimenter is not content with the results of the inferences on the observed data only, but also explores the space delimited by the choice of the models to compare, as well as by the choice of the parameter range (Fig. 10). Compared to a decade ago, nowadays the execution time of an inference pipeline (model comparisons, parameter estimation) has become almost negligible compared to the preliminary steps: identification of the populations to be sampled, sampling campaign, nucleic acid extraction, sequencing, raw data processing. This time saving should benefit the reflection on the scenarios to be explored and compared and on exploring the limits of the inferential approach adopted, notably by relying on the analysis of custom-made simulated data.

## Acknowledgements

C.R has greatly benefited from the support of the I-SITE ULNE foundation (Université Lille Nord-Europe) through the grant “Jeunes Chercheurs 2020” (GAFA project). The simulations were performed on the Core Cluster of the Institut Français de Bioinformatique (IFB) (ANR-11-INBS-0013).

